# Phenotypic and transcriptional response of *Daphnia pulicaria* to the combined effects of temperature and predation

**DOI:** 10.1101/2022.02.24.481768

**Authors:** Aaron Oliver, Hamanda B. Cavalheri, Thiago G. Lima, Natalie T. Jones, Sheila Podell, Daniela Zarate, Eric Allen, Ronald S. Burton, Jonathan B. Shurin

## Abstract

*Daphnia*, an ecologically important zooplankton species in lakes, shows both genetic adaptation and phenotypic plasticity in response to temperature and fish predation, but little is known about the molecular basis of these responses and their potential interactions. We performed a factorial experiment exposing laboratory-propagated *Daphnia pulicaria* clones from two lakes in the Sierra Nevada mountains of California to normal or high temperature (15°C or 25°C) in the presence or absence of fish kairomones, then measured changes in life history and gene expression. Exposure to kairomones increased upper thermal tolerance limits for physiological activity in both clones. Cloned individuals matured at a younger age in response to higher temperature and kairomones, while size at maturity, fecundity and population intrinsic growth were only affected by temperature. At the molecular level, both clones expressed more genes differently in response to temperature than predation, but specific genes involved in metabolic, cellular, and genetic processes responded differently between the two clones. Although gene expression differed more between clones from different lakes than experimental treatments, similar phenotypic responses to predation risk and warming arose from these clone-specific patterns. Our results suggest that phenotypic plasticity responses to temperature and kairomones interact synergistically, with exposure to fish predators increasing the tolerance of *Daphnia pulicaria* to stressful temperatures, and that similar phenotypic responses to temperature and predator cues can be produced by divergent patterns of gene regulation.

## Introduction

Many species are at risk of extinction as the environment changes at an unprecedented pace. Ecological and evolutionary research aims to predict species’ responses to anthropogenic environmental change, such as global warming and introduction of predators [1, 2]. To cope with such stressors, species can either migrate to more suitable habitats or adapt to new conditions. Understanding the potential and limits of genetic adaptation and phenotypic plasticity to maintain fitness is critical in order to predict persistence of populations and changes in biodiversity under environmental stresses [3]. The mechanisms by which organisms are adapted to present day conditions throughout their ranges therefore provide a basis for predicting extinction, persistence, and changes in biodiversity in a rapidly changing environment [4, 5].

Plasticity and genetic adaptation impact organismal fitness in response to environmental change. Plasticity allows a genotype to express multiple phenotypes in different environments whereas genetic adaptation arises from environment-dependent variation in fitness of different genotypes [6]. Evidence for rapid evolution in natural systems, particularly in response to human-induced environmental change, indicates that most of the observed changes are not genetically based, but rather a consequence of plasticity [7]. Plasticity at level of gene expression is one of the most important mechanisms for coping with stress [8, 9], yet the magnitude and genetic basis of such plasticity remains largely unknown.

Plasticity arises from differential gene expression patterns in response to environmental cues [10]. Variable levels of plasticity may evolve in a population if reaction norms differ across genotypes and the slope of the reaction norm is correlated with fitness [11]. Either decreases or increases in phenotypic plasticity could contribute to adaptation to variable environments depending on the rate and predictability of environmental change [6, 12].

Several factors could constrain the evolution of plasticity at the transcriptome level. First, genetic variation potential for plasticity could be limited or absent. Alternatively, even when genetic variation is present, its evolution could be constrained by costs of the mechanisms underlying plastic responses [13]. A trade-off is expected where enhanced plasticity would be beneficial in more spatially or temporally variable environments, but detrimental in a stable environment [14, 15]. In addition, if environmental variation is unpredictable, then plasticity may fail to match phenotypes to the environment to produce higher fitness.

*Daphnia* is an ecologically important genus of crustacean zooplankton in lakes that transfers energy from phytoplankton to fish and invertebrate predators and exerts top-down grazing control on primary production [16]. *Daphnia* shows both genetic adaptation and plasticity at a phenotypic level in tolerance to two important determinants of fitness: temperature and predation. Previous studies have documented that the response of *Daphnia* populations to thermal stress is strongly correlated with environmental conditions and can be either genetic, plastic or both [17–19]. Although these studies show plastic and evolutionary changes, the co-occurrence of other environmental stressors, such as predation, with temperature might interact in natural populations. Fish predation and temperature impose selection on many of the same traits, and in the same direction [18, 20]. For instance, *Daphnia* often mature at smaller sizes and younger ages in warmer water and when fish are present, yet little is known about the molecular basis of this response among populations or their potential interactive effects.

We collected *Daphnia pulicaria* from two lakes in the Sierra Nevada Mountains, California, USA with different thermal conditions to conduct an experiment to evaluate the degree to which life history, thermal tolerance and gene expression are influenced by temperature and predation. We predicted that warm temperatures and exposure to predators should produce similar responses in life history traits, then used transcriptomics to ask whether the same genes underlie the plastic response to fish kairomones and warming, or if the similar phenotypic effects arise from pleiotropic effects where altered gene regulation affects multiple phenotypic traits. Our goal was to ask whether *D. pulicaria’s* plastic response to one selective agent (fish or warming) magnified or dampened the response to the other in terms of both phenotype and gene expression.

## Materials and methods

### Lake sampling

Gardisky (37.955774, -119.251198) and Blue Lakes (38.051164, -119.270342), located in Inyo National Forest, California, USA, were sampled in late August and early September 2017, respectively. In both lakes we measured temperature throughout the water column at each meter using a field probe (YSI Incorporated, Yellow Springs, Ohio, USA). We also collected live zooplankton from the deepest point of the lake using a 30 cm diameter, 63 μm mesh conical net with a 1 m length through the water column, starting 1 m above the lake bottom. These samples were kept cold until returning to the laboratory, where we searched for *Daphnia pulicaria*. When present, 30 *D. pulicaria* females carrying eggs in the brood pouch were separated in 50 ml centrifuge tubes filled with COMBO medium [21]. Each *Daphnia* individual was considered a maternal line. Each maternal line was cultured for at least twelve generations in separate 50 mL tubes filled with COMBO medium under ambient lab conditions. All *D. pulicaria* were fed non-viable cells of the green alga *Nannochloropsis* sp. (Brine Shrimp Direct, Ogden, Utah, USA) at a constant high rate of 24 × 10^6^ cells every two days.

### Experimental design

We randomly picked one clonal line from each lake to propagate. *Daphnia pulicaria* reproduces asexually under benign conditions. Thus, all individuals propagated asexually in the experiment. We started by randomly picking one mature female from each clone. Upon the release of the second clutch, we isolated three neonates that were separated and moved to separate 100 mL containers containing the same media and algae. All individuals were transferred to fresh media and algae three times a week and reared at 15 ± 1°C and photoperiod 12:12 h light/dark. Propagation continued until 100 neonates were collected for each of the clonal lines, with all individuals used for further study born within a 12-hour period. All neonates were placed into 100 ml jars containing COMBO medium at a density of 3 *Daphnia*/jar [21]. Each jar was randomly allocated to one of four treatments: (1) optimum temperature (15°C) without kairomones (fish cues), (2) high temperature (25°C) without fish cues, (3) optimum temperature with fish cues, and (4) high temperature with fish cues. An optimum temperature near 15°C is consistent with previous reports of thermal preference in *Daphnia pulicaria* [24], although there is variation in tolerable temperatures between *D. pulicaria* genotypes [25]. All *Daphnia* were transferred to fresh medium, with algae (and kairomones in the fish cue treatment) daily. We monitored jars daily for maturation (i.e., release of first clutch into the brood chamber). Upon reaching maturity, individuals were either preserved in RNAlater (Qiagen, USA) and kept in -20°C for subsequent RNAseq analyses or assigned to phenotypic assay (see below). Since RNA was extracted from whole individuals, no food was added during the last 12 hours before sampling to minimize algal RNA contamination (most of which will be digested and hence degraded after 12 hours). The period without food was kept short to minimize starvation-dependent gene regulation.

### Kairomone collection

COMBO medium conditioned by the presence of planktivorous fish was collected daily from a tank containing 5 juvenile rainbow trout (*Oncorhynchus mykiss*; ∼ 5 cm in total length) in 72 L of COMBO. Each day, media containing fish chemical cues was filtered through 0.7 μm mesh membrane filters and added at a concentration of 0.007 fish/L to the predator treatments. This concentration of kairomone media is above the threshold used by other studies to elicit a phenotypic response, such as the concentration of 5 µl of kairomone media per 100 ml of solution [26].

*D. pulicaria* predation by *O. mykiss* in artificially stocked lakes is well established, specifically predation by juvenile rainbow trout [27, 28], *O. mykiss* is one of the most abundant species for stocking previously fishless lakes in the Sierra Nevada mountains, the source of our *Daphnia* samples [29]. *O. mykiss* were fed TetraMin Tropical Flakes (Tetra, Blacksburg, Virginia, USA) prior to kairomone collection, rather than juvenile conspecific daphniids as in some other studies [30, 31]. We selected this method to minimize the presence of conspecific alarm cues in the isolated kairomone media, so that we could measure the phenotypic and genetic responses to only predator-released, diet agnostic kairomone cues. Such research strategies are supported by Mitchell et al. [32], who call for further research to determine the relationship between predator odor and diet-based cues in aquatic systems. Following their nomenclature recommendations, we define the kairomones used in this study to be predator odor cues, as opposed to alarm cues released by injured conspecifics or diet cues released by predators after digesting conspecifics. All procedures involving animals were reviewed and approved by the Institutional Animal Care and Use Committee of the University of California San Diego (IACUC protocol number: S14140).

### Phenotype assays

Individuals from the experimental generation assigned to the phenotypic assay were scored for the following life-history variables: age and size at maturity, and age and number of offspring from the first three clutches. These data were used to calculate intrinsic population growth rate for each maternal line following the Lotka–Euler equation [33]. We also measured critical maximum temperature (CT_max_) using a heat ramping assay. After a 30-minute resting period at ambient temperature, *Daphnia* from the four treatments were transferred to 0.5ml Eppendorf tubes and placed in a thermal heater (4x6 thermal heater, Corning Digital Dry Bath Heater Dual Block). The water temperature increased 0.1°C every 20 seconds. We continuously monitored the state of individuals and recorded the temperature when each individual *Daphnia* lost swimming ability and sank to the bottom of the tube. The temperature when *Daphnia* became immobilized was used as a proxy for CT_max_ [17]. We measured life history traits of 11 to 13 individuals per treatment per lake and CT_max_ of 10 to 12 individuals per treatment per lake. In total, we scored phenotypes of 186 individuals.

### Statistical analysis for phenotype

We log-transformed all continuous variables, except intrinsic growth rate, after visually assessing the probability distribution that best fit the data. We analyzed the effects of the treatments using generalized linear models for each trait using the statistical software R [34]. Temperature treatment, fish cue treatment, source lake and the interactions of these variables were modeled as fixed effects for age and size at maturity, average number of offspring, intrinsic growth rate and CT_max_. Since we expected that CT_max_ would be influenced by body size we included an additive effect of size.

Additionally, to test the effects of slopes of the reaction norms, we calculated the pairwise differences for each dependent variable between each treatment combination within a lake. We computed pairwise differences using the least square mean values based on the generalized linear models for each trait using *lsmeans* function in the *lmerTest* R package [35]. The pairwise difference test was corrected using Tukey’s adjustment for multiple comparisons [36].

### RNA isolation, library preparation and sequencing

We extracted RNA using the TRI Reagent Protocol (Sigma-Aldrich, USA) from samples preserved in RNAlater (Qiagen, USA). Any remaining genomic DNA was removed using the TURBO DNA-free Kit (Invitrogen, USA). Each treatment included five biological replicates, with each replicate comprised of 35 clonal *Daphnia* individuals. Extracted RNA was stored in -80°C until sequencing. Quality of the isolated RNA was assessed using RNA Nano 6000 Assay Kit of the Agilent Bioanalyzer 2100 system (Agilent Technologies, CA, USA) and mRNA-seq libraries were constructed using NEBNext UltraTM RNA Library Prep Kit for Ilumina (NEB, USA) following manufacturer’s recommendations. A total amount of 1 μg RNA per replicate was used as input material for RNA-seq library preparations. Samples that did not reach 1 μg RNA were pooled with a biological replicate grown under the same conditions, a process described by Takele Assefa et al. [37], leaving 18 replicates from Blue Lake and 16 replicates from Gardisky Lake. Index codes were added to attribute sequences to each sample and all 34 libraries were clustered on a cBot System using the PE Cluster kit cBot-HS (Illumina). After generating the clusters, libraries were sequenced using the Illumina HiSeq 2000 platform and 150 bp paired-end reads were generated.

### Read cleaning and transcriptome comparison

The raw reads were preprocessed using fastp v. 0.23.1 [38] with the command line options “-x -f 18 -F 18”. These options enabled polyX tail trimming and removed 18 low-quality bases from the beginning of all forward and reverse reads, respectively. To check for contamination in our read set, we used Kraken2 v. 2.0.9 [39] with a taxonomy database constructed from the NCBI nr database [40] as of August 2021. A random subset of 1,000,000 read pairs was used for each Kraken2 run per sample. All downstream analyses were based on the remaining cleaned, validated data.

Separate *de novo* assemblies for were performed on the Blue and Gardisky samples using Trinity v. r20140413p1 [41] with parameter min_kmer_cov set to 2. Transcriptome completeness was assessed using BUSCO v. 5.2.2 [42] with the arthopoda_odb10 lineage dataset, and also by comparison to the following published, annotated genomes: *D. pulicaria* LK16 [43], *D. pulex* TCO [44], *D. galeata* M5 [45], *D. magna* SK [46] and *Eulimnadia texana* JT4(4)5-L [47].

### Differential expression analysis

Relative expression levels of annotated genes were estimated by independently mapping RNA-Seq reads from each lake sample back to the annotated *D. pulicaria* LK16 gene set using STAR v. 2.7.9a [48]. A read count matrix was generated from these alignments using featureCounts v. 2.0.1 [49]. Genes with read counts below 10 across all conditions were filtered from the remaining pipeline. The significance of difference in gene expression of each treatment for each genotype was determined with the R package DESeq2 v. 1.32.0 [50], using a model based on negative binomial distribution. Statistical significance (p-value) was adjusted using the q-value obtained from the false discovery rate [51], with a q-value < 0.05 and |log2(foldchange)| > 1 set as the threshold for significant differential expression. Counts of differentially expressed genes across treatments were visualized using the R package VennDiagram v. 1.7.1 [52]. A principal component analysis (PCA) was visualized using read counts corrected through the variance stabilizing transformation [53] function of DESeq2, The PCA was generated using corrected read counts from every gene, rather than the DESeq2 default of using only the 500 most variable genes. These correct read counts were also used to perform a PERMANOVA test [54] through the R package vegan v. 2.5-7 [55] with 999 permutations. The groups tested with PERMANOVA were based on lake, temperature, fish cues, and their interactions terms.

Gene Ontology (GO) enrichment analyses were performed to determine whether significantly differentially expressed gene sets were enriched for any biologically relevant functional categories [56]. GO term enrichment lists were generated using GOseq v. 1.44 [57] for the following one-factor comparisons: Blue, temperature rise with kairomones; Blue, temperature rise without kairomones; Gardisky, temperature rise with kairomones; Gardisky, temperature rise without kairomones. Further control of the False Discovery Rate was performed using the Benjamini–Hochberg method [58], and the cutoff for enriched GO terms was an adjusted p-value less than 0.05.

### Orthology of differentially expressed genes

*Daphnia pulex* genes previously demonstrated to be differentially expressed in response to temperature and/or predator kairomones were accessed from the *Daphnia* Stressor Database [59] as of September 2021. Predicted protein sequences for these genes were downloaded from wFleaBase [60]. These genes were compared to the annotated *Daphnia pulicaria* LK16 genome using OrthoVenn2 [61] with an e-value of 1e-5. OrthoVenn2 results were visualized using the R package VennDiagram v. 1.7.1 [52].

## Results

### Phenotype

Several traits in the two clones responded differently to temperature and fish cues (Table 1). The Gardisky Lake clone matured at a younger age than Blue Lake (Lake: p < 0.001, Table 1), but age at maturity was earlier for both clones in the presence of fish cues at high temperature (Temp x Fish: p-value = 0.006, Table 1, Fig 1). The post-hoc test (least square means) showed that fish had an effect only on the Blue Lake clone reared at 25°C. At 25°C, Blue Lake replicates reared with fish cues matured 0.55 ± 0.16 days earlier compared to replicates without fish cues (Fig 1).

**Fig 1.**
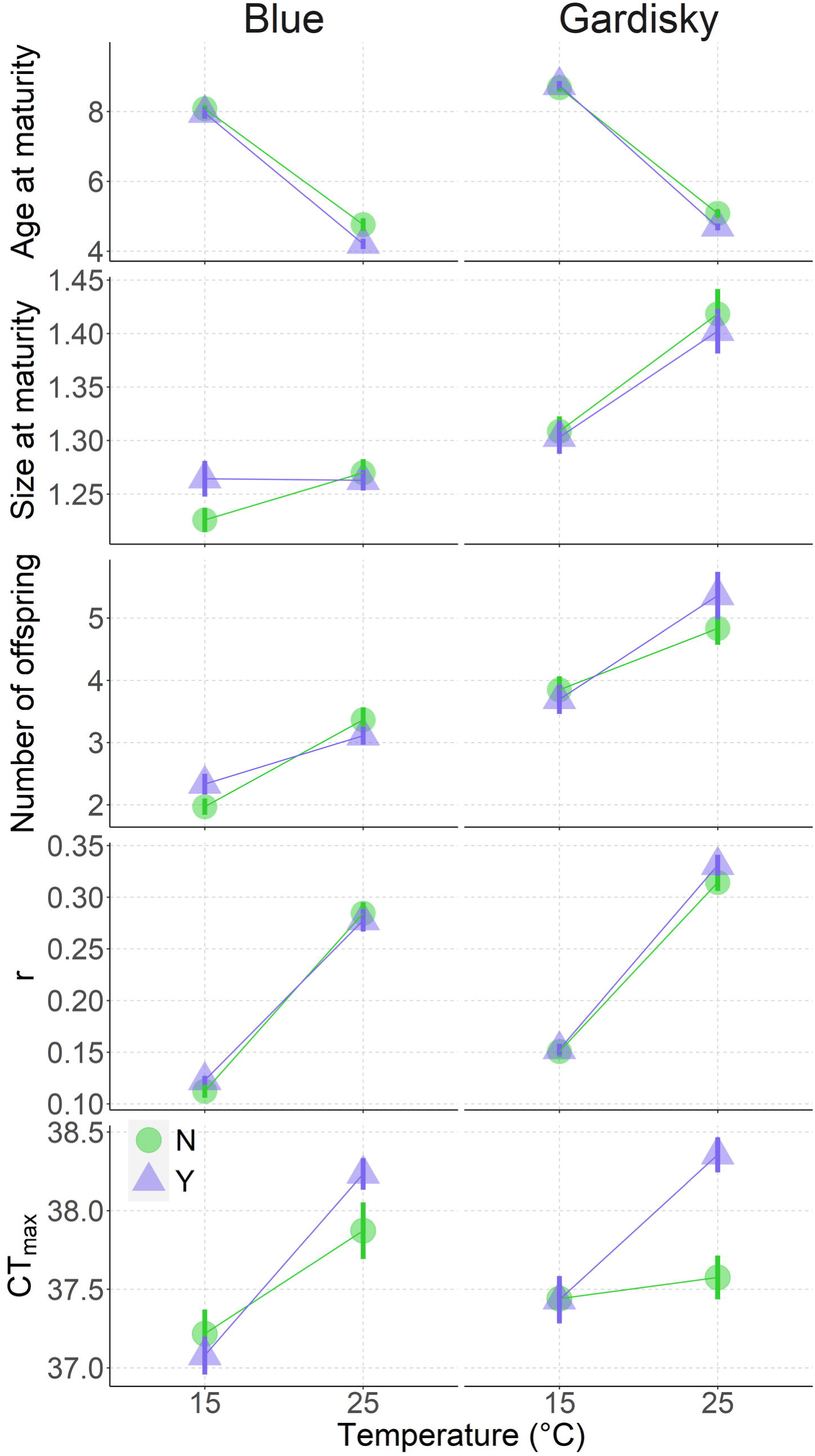
Predation cues influence critical maximum temperature of Daphnia pulicaria. Means ± 1 S.E.M. of age at maturity (days), size at maturity (mm), average number of offspring of the first three clutches, intrinsic growth rate (r), and critical maximum temperature (CTmax, °C) of Daphnia pulicaria collected at Blue and Gardisky Lakes during summer 2017 in response to temperature (15°C and 25°C, x-axis) and predation cues treatments. Circles show response without fish cues and triangles response to fish cues.

**Table 1.** Results of the generalized linear model analysis of the life-history traits of *Daphnia pulicaria*. Clonal lines were collected from Blue and Gardisky Lakes and reared under different temperature (15°C or 25°C) and fish cue treatments (presence or absence).

| Factor | Df | Deviance | Deviance residuals | Pr (>Chi) |
| --- | --- | --- | --- | --- |
| **Age at maturity** |  |  |  |  |
| Temperature | 1, 184 | 16.0918 | 3.1609 | **< 0.001** |
| Fish cues | 1, 183 | 0.1340 | 3.0270 | **0.002** |
| Lake | 1, 182 | 0.3961 | 2.6309 | **< 0.001** |
| Temp x Fish | 1, 181 | 0.1036 | 2.5273 | **0.006** |
| Temp x Lake | 1, 180 | 0.0020 | 2.5253 | 0.7098 |
| Fish x Lake | 1, 179 | 0.0147 | 2.5106 | 0.3074 |
| Temp x Fish x Lake | 1, 178 | 0.0011 | 2.5096 | 0.7800 |
| **Size at maturity** |  |  |  |  |
| Temperature | 1, 182 | 0.1078 | 0.9068 | **< 0.001** |
| Fish cues | 1, 181 | 0.00002 | 0.9068 | 0.9349 |
| Lake | 1, 180 | 0.2726 | 0.6341 | **< 0.001** |
| Temp x Fish | 1, 179 | 0.0054 | 0.6286 | 0.1989 |
| Temp x Lake | 1, 178 | 0.0382 | 0.5904 | **< 0.001** |
| Fish x Lake | 1, 177 | 0.0046 | 0.5858 | 0.2368 |
| Temp x Fish x Lake | 1, 176 | 0.0024 | 0.5833 | 0.3917 |
| **Average number of offspring** |  |  |  |  |
| Temperature | 1, 87 | 2.7408 | 9.7072 | **< 0.001** |
| Fish cues | 1, 86 | 0.0403 | 9.6669 | 0.3436 |
| Lake | 1, 85 | 5.7215 | 3.9454 | **< 0.001** |
| Temp x Fish | 1, 84 | 0.0105 | 3.9349 | 0.6290 |
| Temp x Lake | 1, 83 | 0.0826 | 3.8524 | 0.1751 |
| Fish x Lake | 1, 82 | 0.0030 | 3.8494 | 0.7963 |
| Temp x Fish x Lake | 1, 81 | 0.2120 | 3.6374 | **0.0297** |
| **Intrinsic growth rate** |  |  |  |  |
| Temperature | 1, 87 | 0.6194 | 0.0962 | **< 0.001** |
| Fish cues | 1, 86 | 0.0007 | 0.0954 | 0.3087 |
| Lake | 1, 85 | 0.0327 | 0.0627 | **< 0.001** |
| Temp x Fish | 1, 84 | 0.0000 | 0.0627 | 0.9713 |
| Temp x Lake | 1, 83 | 0.0003 | 0.0624 | 0.5032 |
| Fish x Lake | 1, 82 | 0.0003 | 0.0620 | 0.4853 |
| Temp x Fish x Lake | 1, 81 | 0.0014 | 0.0606 | 0.1665 |
| **Critical maximum temperature (CT<sub>max</sub>)** |  |  |  |  |
| Size at maturity | 1, 91 | 0.0003 | 0.0256 | 0.1655 |
| Temperature | 1, 90 | 0.0082 | 0.0173 | **< 0.001** |
| Fish cues | 1, 89 | 0.0009 | 0.0163 | **0.0125** |
| Lake | 1, 88 | 0.0004 | 0.0159 | 0.1112 |
| Temp x Fish | 1, 87 | 0.0015 | 0.0144 | **0.0019** |
| Temp x Lake | 1, 86 | 0.0006 | 0.0138 | 0.0523 |
| Fish x Lake | 1, 85 | 0.0003 | 0.0134 | 0.1356 |
| Temp x Fish x Lake | 1, 84 | 0.0000 | 0.0134 | 0.5433 |

Temperature affected size at maturity differently in the two populations (Temp x Lake: p < 0.001, Table 1) but fish cues had no main effect across clones (Fish: p = 0.093, Table 1, Fig 1). Post hoc tests revealed that the Blue Lake clone matured at 1.26 ± 0.006 mm regardless of temperature, while Gardisky matured at 15°C with 1.31 ± 0.01 mm compared to 1.41 ± 0.01 mm at 25°C.

The average number of offspring for the first three clutches showed a significant three-way interaction (Temp x Fish x Lake: p = 0.029, Table 1, Fig 1). For the Blue Lake clone, offspring number only varied with temperature; on average Blue had 1.1 ± 0.11 more offspring at 25°C compared to 15°C. In contrast, the effect of temperature on fecundity was greater for the Gardisky Lake clone when fish cues were present. Individuals had 1.67 ± 0.30 more offspring at 25°C than 15°C in the presence of fish cues, while without fish cues the difference between 25°C and 15°C response was 0.98 ± 0.23 offspring.

The intrinsic growth rate was significantly affected by the main effects of temperature and lake (Temp: p < 0.001 and Lake: p < 0.001, Table 1, Fig 1). At each temperature, the Gardisky Lake clone had higher intrinsic growth rates compared to the Blue Lake clone. For instance, the Gardisky clone had an *r* = 0.15 ± 0.001*day^-1^ at 15°C, while Blue had *r* = 0.11 ± 0.004*day^-1^. At 25°C, the intrinsic growth rate was 0.32 ± 0.006*day^-1^ for the Gardisky Lake clone, while the growth rate for the Blue Lake clone was 0.28 ± 0.007*day^-1^.

CT_max_ increased significantly when *Daphnia* were reared at 25°C compared to 15°C. CT_max_ was significantly affected by the interaction between temperature and fish cues (Temp x Fish: p = 0.002, Table 1) and showed a marginally significant interaction between temperature and lake (Temp x Lake: p = 0.052, Table 1), but no main effect of lake. Post hoc tests showed that at 15°C, fish cues had no effect on CT_max_. The thermal tolerance of the Gardisky clone increased by 0.8 ± 0.12°C in the presence of fish cues, while the Blue Lake clone increased by 0.3 ± 0.14°C at 25°C.

### Gene expression

For the Blue Lake clone, sequencing produced on average 42,306,315 raw pair-ended reads for each sample. Across all Blue Lake samples, 79.4% of the cleaned reads (after removing adaptor sequences, low-quality and ambiguous sequences) were mapped to the D. pulicaria LK16 primary gene set. The Gardisky clone produced on average 44,025,859 raw pair-ended reads for each sample and 80.0% of the cleaned reads mapped to the D. pulicaria LK16 primary gene set. A breakdown of read mapping rate and estimated read set contamination by sample is visualized in S1 Fig. *De novo* assemblies of the Blue and Gardisky clones show sequence similarity to previously annotated *Daphnia* genomes (S2 Fig) and high levels of completeness consistent with previously annotated arthropod genomes (S3 Fig). A PCA plot generated using DESeq2-corrected read counts (Fig 2) shows clear divergence in the gene expression between clones under all growth conditions. A PERMANOVA test of gene expression data shows that gene expression was affected by both source lake (Pseudo F = 9.6582, R^2^ = 0.22284, p = 0.001) and temperature treatments (Pseudo F = 4.367, R^2^ = 0.10078, p = 0.001), but gene expression was not significantly different between predation treatments (Pseudo F = 0.6706, R^2^ = 0.01547, p = 0.746).

**Fig 2.**
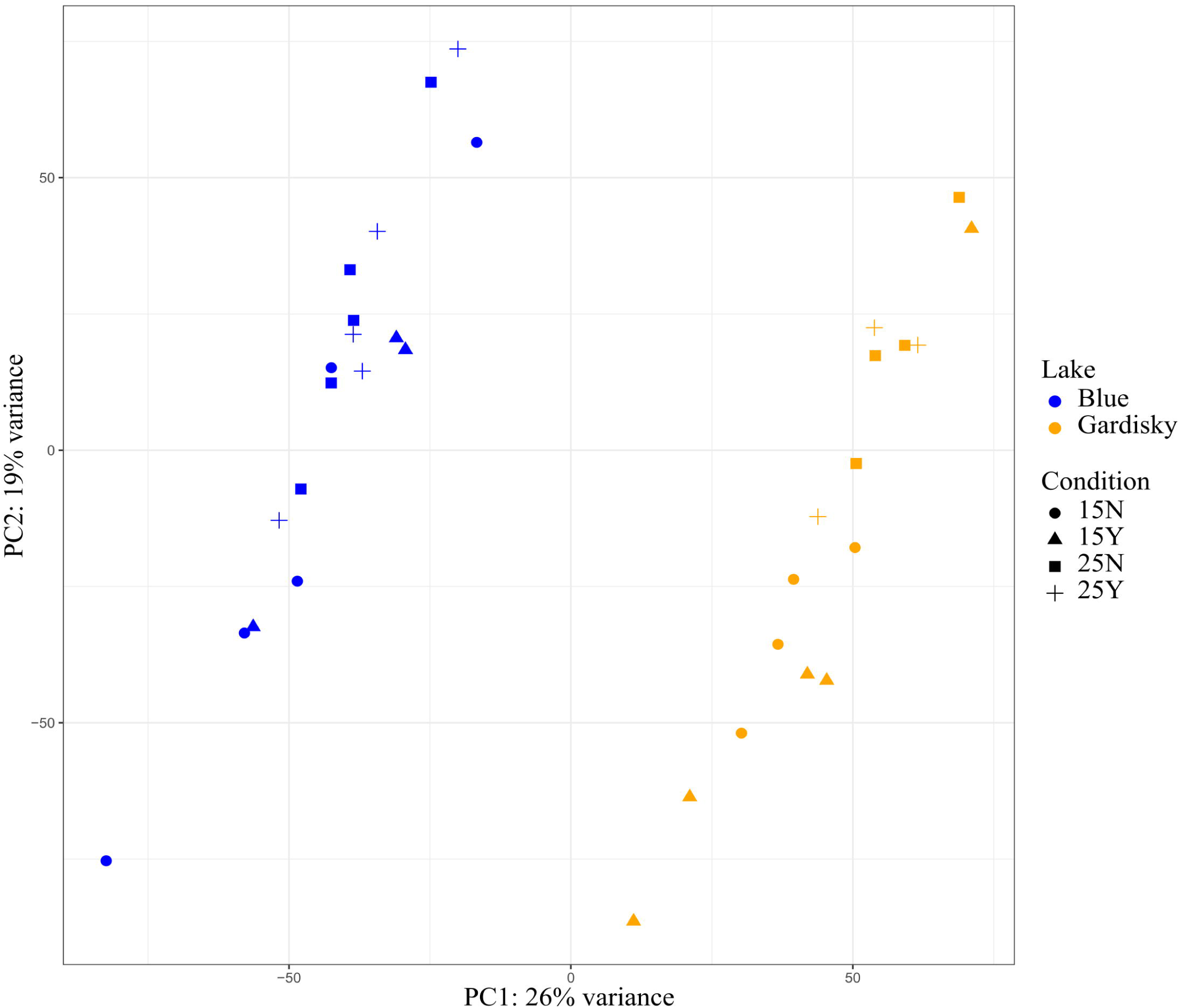
Genotype, not treatment group, is the dominant source of differential gene expression. Principal component analysis of gene expression patterns generated through DESeq2 prominently separates individuals by source lake along the first principal component. Temperature treatments (25N vs. 15N and 25Y vs. 15Y) produced the greatest number of differentially expressed genes (DEGs) for both the Blue and Gardisky clones (Fig 3). A total of 320 genes were differentially expressed between one-factor comparisons for the Blue Lake clone: 95 due to temperature differences with predation cues, 292 due to temperature differences without predation cues, and 4 due to fish cues at 25°C (Fig 3a). For the Gardisky Lake clone, we identified 575 DEGs. In total, 396 DEGs were specific to 25N vs. 15N and 52 DEGs were specific to 25Y vs. 15Y. These two treatment comparisons shared 127 genes (Fig 3b). Predator cues alone had no measurable effect on gene expression when comparing between Gardisky clones grown under the same temperature. Tables listing all differentially expressed genes are presented in Supplementary Tables 1a and 1b.

**Fig 3.**
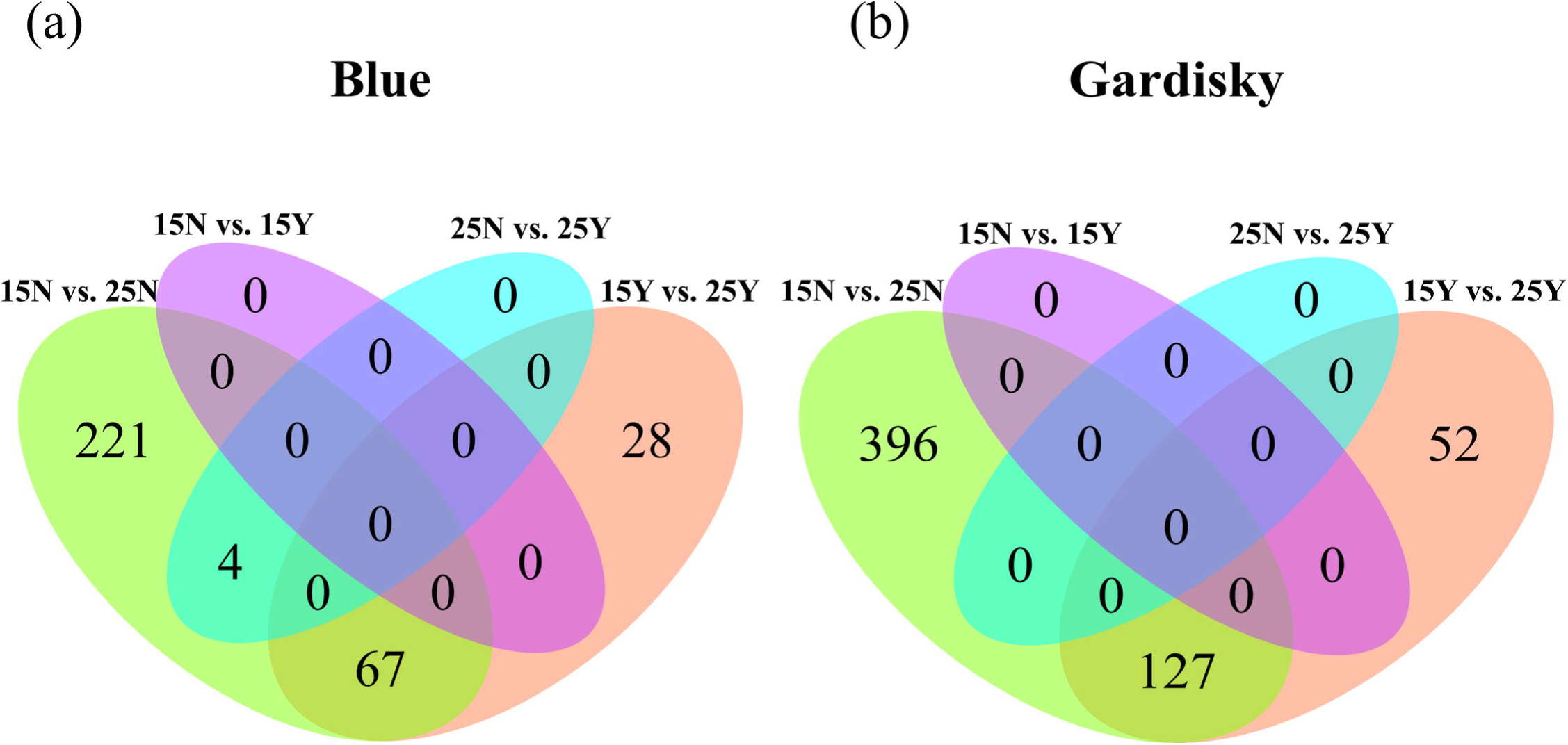
Many genes are differentially expressed in response to temperature changes, while kairomones have minimal observable effect at the level of transcript expression. Venn diagram of differentially expressed genes for *Daphnia pulicaria* from **(a)** Blue and **(b)** Gardisky Lakes in response to temperature and predation cues. Cyan and purple sections illustrate differential expression from the introduction of predator cues (Y vs. N for the presence and absence of fish cues), while the green and orange sections illustrate differential expression due to temperature change. Treatment abbreviations: 15Y, 15°C with kairomones; 15N, 15°C without kairomones; 25Y, 25°C with kairomones; 25N, 25°C without kairomones.

Comparisons of significant GO terms for differentially expressed genes were difficult to interpret due to the limited specificity of the functional information they provide. Genes related to proteolysis, chitin turnover, and vision appeared to be upregulated at higher water temperatures, but no significantly over- or under-represented GO terms were detected in response to kairomones. GO terms for phototransduction and visual perception were enriched in all temperature comparisons, while protease and peptidase activity were enriched in every comparison but 25Y vs. 15Y in the Blue clone. Meanwhile, terms associated with chitin binding were enriched in every comparison but 25Y vs. 15Y in the Gardisky clone. Complete lists of all enriched GO terms are given as Supplementary Tables 2a-2d.

### Gene orthology

From the Daphnia Stressor Database, we retrieved 386 genes that responded to kairomones in *D. pulex* across 7 studies, and 2660 genes that responded to temperature changes in *D. pulex* across 4 studies. 315 of these previously documented kairomone-responsive genes (81.6%) have orthologs in the *D. pulicaria* LK16 genome, while 1796 of the temperature-responsive genes (67.5%) are orthologous to genes in *D. pulicaria* (Fig 4a). 35% of the temperature responsive *D. pulicaria* DEGs in this study are orthologous to previously reported temperature-responsive genes from *D. pulex*, but only one of the four genes in the Blue clone that responded to kairomones was orthologous to a previously described kairomone-sensitive gene (Fig 4b). On the other hand, we observed that 18% of described kairomone-responsive genes in *D. pulex* have orthologs that respond to temperature in *D. pulicaria*, consistent with earlier phenotypic observations (Fig 1) showing an interactive relationship between these variables. Lists of gene identifiers for orthologous clusters and singleton genes are available in Supplementary Files 1 and 2.

**Fig 4.**
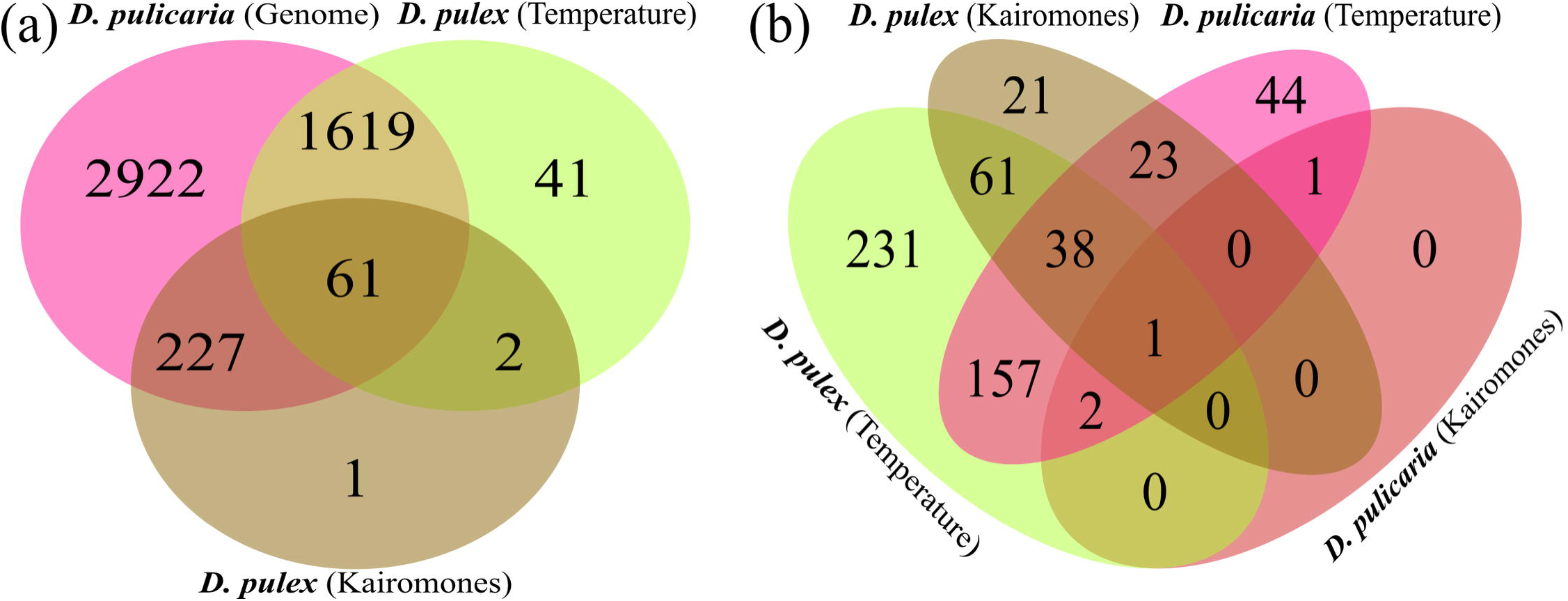
Gene orthology analysis reveals similarities between stress-responsive genes in *D. pulicaria and D. pulex*. Venn diagrams of orthologous gene clusters between *D. pulex* DEGs from the *Daphnia* Stressor Database and **(a)** the *D. pulicaria* genome or **(b)** DEGs from *Daphnia pulicaria.* Numbers indicate protein families with at least two members, excluding “singletons” that have no orthologous or paralogous matches. Gene identifiers for orthologous clusters and singleton genes are available in **(a)** Supplementary File 1 and **(b)** Supplementary File 2.

## Discussion

*Daphnia* clonal lines that were stressed by temperature and/or predator cues exhibited similar phenotypic responses associated with distinctly different patterns of gene regulation. Temperature was the most important factor driving changes in gene expression, as most of the DEGs in both clones responded to this variable, versus only a handful to kairomone cues. Yet fish cues magnified the effect of 10°C of warming on age of maturity and increased the critical maximum temperature at which *D. pulicaria* lose motor control when individuals were reared at high temperatures, but not at their optimum temperature. Genetic pathways that underlie the plastic response to predators and temperature may be subject to different regulatory pressures, but phenotypic responses suggest that exposure to predator cues may increase thermal tolerance and affect life history response to warming.

### Phenotypic responses

Both clones matured at a younger age and produced more offspring when reared in high temperature, thus having a higher intrinsic growth rate compared to lower temperature conditions (Fig 1). Indeed, high temperature increases metabolic rate, often leading to a larger reproductive investment and faster development [62, 63]. In addition, we found an interactive effect of temperature and predation on age at maturity and critical maximum temperature.

Our finding that individuals reared with predator cues had higher thermal tolerance and matured younger at 25°C than those reared without predator cues agree with other studies that also found synergistic effect of these two stressors in *Daphnia* and other aquatic organisms [64–69]. In *D. pulex*, after seven generations, Tseng and O’Connor observed increased thermal plasticity in individuals reared in higher temperatures only when they were also reared with predators [64]. Zhang et al. tested resurrected *D. magna* population from time periods without fish and from a high-density fish period [65]. They found that *Daphnia* that coexisted with fish exhibited earlier maturation in high temperatures compared to *Daphnia* from the pre-fish period. Earlier maturation under predation risk is common in size-dependent predation of *Daphnia* where predators, such as fish, prefer larger prey items, inducing a smaller size at maturity and earlier age of first reproduction [63]. Our results agree with these studies showing that fish cues magnify the life history response to temperature and show that exposure to fish predation can increase the tolerance of stressfully high temperatures.

### Genetic responses to temperature

We observed upregulation of serine peptidases in response to high temperature treatment for both clones. These enzymes are the most important digestive proteases in *D. magna* [70]. They are involved in proteolysis in the digestive system, the process by which peptide bonds in proteins are broken generating free amino acids. This upregulation may indicate a necessity to accommodate higher feeding rates caused by increased metabolism in high temperatures [62, 63]. Moreover, the *D. pulex* genome contains many peptidase genes, which might indicate adaptation to high variation in food availability in aquatic environment [71].

Genes related to visual perception were also upregulated in response to high temperature treatment in both clones. Vision has previously been associated with vertical migration, because visible light is a proxy for both *Daphnia’s* visibility to predators and the amount of incoming photodamage due to UV radiation at near-surface depths [72].

Vertical migration can serve as either a predator avoidance mechanism induced directly from kairomones or a diel behavior triggered by temperature change [72]. Higher temperatures can serve as a proxy for shallower depths, and therefore more incoming radiation. This may explain why our dataset shows enrichment of phototransduction genes as a response to temperature while previous studies have observed phototransduction gene enrichment in response to kairomones [31].

### The missing kairomone response: behavioral changes and proximate cues

In total, we only detected 4 DEGs across two *Daphnia pulicaria* clones in response to kairomones from *Oncorhynchus mykiss.* Such a low number of DEGs related to kairomone exposure is unexpected, but not unprecedented in previous *Daphnia* studies. One study on *Daphnia galeata* M6 by Tams et al. also found 4 differentially expressed transcripts in response to kairomones from the fish *Leuciscus idus* using a two-factor analysis [73], and another study found no DEGs in *Daphnia magna* Iinb1 when exposed to kairomones from stickleback fish [74]. Tams et al. hypothesize that life history changes might be associated with only a few genes compared to morphological defenses, or that regulation could be post-translational and therefore would not appear in an RNA sequencing study [73]. Defense strategies can also include behavioral changes such as vertical migration [75, 76] and the transition into a state of alertness [77], which RNA expression data may not fully capture. In contrast, morphological changes such as neckteeth formation have been associated with far more genes, such as 230 DEGs found in *D. pulex* in response to *Chaoborus* kairomones [78].

Miehls et al. find that the spiny water flea *Bythotrephes longimanus* displays no changes in life history or morphology when exposed to kairomones from the predator *Perca flavescens,* but increased temperature does induce morphologic defense against predation [79]. The authors hypothesize that temperature is a stronger predictor of predation risk than kairomones. This is because adult *P. flavescens* are present year-round but only juveniles are gape-limited in their consumption of *B*. *longimanus,* such that the morphologic response of increased tail length is only effective in discouraging predation from juveniles and not adult fish. If there is no difference between the kairomones released from juvenile and adult *P. flavescens,* then temperature could serve as a proximate cue for predation risk that could be damped through some phenotypic response.

The interactions between *B*. *longimanus* and *P. flavescens* closely mirror predation of *D. pulicaria* by *O. mykiss*, as *O. mykiss* are among the most abundant stocked fished in the Sierra Nevada lakes [29] and thus *D. pulicaria* are exposed to their kairomones year-round. Juvenile *O. mykiss* prey on *Daphnia* as a large percentage of their diet, while trout larger than 350 mm are primarily piscivorous [27]. A morphological defense response targeted solely at juvenile trout might require either a juvenile-specific odor, a dynamic response based on cue concentration, or some additional proxy for *O. mykiss* life stage that would suggest a high concentration of juveniles. Riessen and Gilbert propose that *D. pulex* may use water temperature to serve as a proximate cue for predator densities [80], so water temperature as a proxy for predator life stage is not implausible. Our dataset found some overlap between kairomone-responsive genes in *D. pulex* and temperature-responsive genes in *D. pulicaria,* so an intermediate defensive response may be inducible through heat alone as found by Suppa et al. [81].

However, behavior-based predator avoidance strategies may not require such specificity. *O. mykiss* can produce the kairomone 5α-cyprinol sulfate in their bile without consuming *Daphnia*, and this compound induces diel vertical migration in *D. pulex* [75]. Vertical migration still has an associated tradeoff in fitness [82], but this defense might be preferred when food sources are abundant and temperature gradients are tolerable [83]. Another study found that *D. pulicaria* lineages that coexist with *O. mykiss* vertically migrate without kairomone exposure, while *D. pulicaria* lineages without prior contact with *O. mykiss* require kairomone exposure to induce diel vertical migration [84]. Thus, some environments with high predation might select for *Daphnia* genotypes that respond to predation primarily through behavioral responses, such as vertical migration.

### The missing kairomone response: predator- and diet-specific cues

An alternative explanation for the lack of DEGs may lie in the subtle differences between kairomone sources. Some kairomones are innately released by the predator (predator odors), from damaged prey (alarm cues), and from digested prey excreted by the predator (dietary cues). This study and Tams et al. explicitly avoided the addition of conspecific alarm cues in our kairomone mixture as to measure only the effect of predator odors [73]. Other studies, such as [30, 31], tested responses to a mixture of predator-specific odors and alarm cues released by consumed conspecifics. Mitchell et al. call for the separation of nomenclature between cues released from the predator regardless of diet and diet-specific cues [32], so that researchers can more accurately discern the independent effects of these different kairomone compounds.

Previous research on the interaction between predator odors and alarm cues is mixed. Some studies propose that defense systems rely on both types of compounds to induce measurable defense responses [85, 86], while others observe that alarm cues by themselves can induce separate responses than predator odors alone [87, 88]. Alarm cues are thought to be nonspecific, as *Daphnia* respond differently to different predators [89] and alarm cues alone might be unable to induce defense responses effective against a specific predator [90]. The most effective defense responses may therefore require predator odors to identify the type of predator combined with alarm cues to induce an immediate response. A required mixture of predator odors and alarm cues for maximum observed response in *D. pulicaria* would succinctly explain why we observed such a minor response in our experiment. Results from this study support the conclusion that the detection of predator odors alone may be insufficient to induce defense responses at the level of gene expression, at least in the context of *D. pulicaria – O. mykiss* interactions.

## Conclusion

Our study highlights that expression patterns of genes differed between *Daphnia pulicaria* clones while phenotypic responses and interactions were qualitatively similar, suggesting that different transcriptomic responses can result in similar phenotypes. Other studies have found distinct gene expression patterns among individuals from different zooplankton species [91, 92]. These results suggest that diverse genetic pathways can give rise to similar phenotypic plastic responses to environmental stress. We also showed synergistic interactions between temperature and predation for some traits. Our results show that *Daphnia* can have similar phenotypes through distinct molecular mechanisms and the synergistic effects of temperature and predation may represent an important mechanism for organisms to adapt to a rapidly changing environment. As one of the first studies incorporating the *Daphnia pulicaria* genome, we hope to lay the groundwork for future differential expression studies through incorporation into knowledge bases such as the *Daphnia* Stressor Database. Overall, we find that the interactions between stressors are heavily genotype dependent, and echo the sentiment that novel research strategies are necessary [32] to further investigate the chemical signals of predation in *Daphnia*.

## Data Availability

Read sets generated and/or analyzed during the current study are available on the Sequence Read Archive (SRA) under BioProject PRJNA698341. The SRA Accession numbers for the individual paired sets of reads associated with Blue Lake are SRR13591461, SRR13591462, SRR13591463, SRR13591464, SRR13591465, SRR13591466, SRR13591471, SRR13591482, SRR13591485, SRR13591486, SRR13591487, SRR13591488, SRR13591489, SRR13591490, SRR13591491, SRR13591492, SRR13591493, and SRR13591494, The SRA Accession numbers for the individual paired sets of reads associated with Gardisky Lake are SRR13591467, SRR13591479, SRR13591480, SRR13591481, SRR13591483, and SRR13591484. Phenotype data, used to generated Fig 1, is included as S3 File.

## Acknowledgements

We are grateful to the following people for their assistance during the execution of this study: Jennifer Leong, Didra Felix, Carol Blanchette, Brent Salzmann, Kim Rose, Scott Forster, and Josh Kohn. Funding was provided by National Science Foundation DEB grant (1457737) to JBS, Brazilian Federal Agency CAPES (13768-13-1) graduate scholarship to HBC and research funding by Frontiers of Innovation Scholars Program of the University of California San Diego (3-G3056) to HBC. The funders had no role in study design, data collection and analysis, decision to publish, or preparation of the manuscript. The work was performed in part at the University of California Valentine Eastern Sierra Reserve.

## Supplementary Tables, Figures, and Files

**S1 Tables 1a-1b. Data associated with Fig 3.** Differentially expressed genes in the **(a)** Blue Lake clone and **(b)** the Gardisky Lake clone. The comparison column denotes if genes were responsive to temperature and/or kairomone treatments.

**S2 Tables 2a-2d. Gene Ontology of differentially expressed genes.** Upregulated Gene Ontology terms for comparisons between: **(a)** Blue Lake clone, 15Y and 25Y treatments, **(b)** Blue Lake clone, 15N and 25N treatments, **(c)** Gardisky Lake clone, 15Y and 25Y treatments, and **(d)** Gardisky Lake clone, 15N and 25N treatments.

**S1 Fig. Read-based taxonomy assignment consistent with mapping rates to the *Daphnia pulicaria* genome.** Panel **(a)** shows Kraken2 read taxonomy on a random subset of one million cleaned reads from each sample. Panel **(b)** plots the alignment rate of the cleaned reads from each sample to the chromosomes of *Daphnia pulicaria* LK16 using the STAR aligner.

**S2 Fig. Blue and Gardisky transcripts are highly similar to *Daphnia pulicaria* LK16.** Results of a two-way average nucleotide identity (ANI) matrix and associated dendrogram for *the de novo* assembled transcriptomes from Blue and Gardisky Lakes, predicted genes from *Daphnia* species with annotated genomes, and the annotated gene set of the related arthropod *Eulimnadia texana*. Our transcriptome sets are closest to those from *D. pulicaria*, within the well-described *Daphnia pulex-pulicaria* species complex.

**S3 Fig. Blue and Gardisky transcriptomes are highly complete and highly duplicated.** Bar graph of gene set completeness as determined by BUSCO using the arthropod marker gene set. Our *de novo* assembled transcriptomes have completeness scores comparable to or better than genome-derived transcript sets from other *Daphnia* studies. However, Blue and Gardisky transcriptome sets have additional gene duplication in our transcriptome due to the inclusion of gene isoforms from *de novo* assembly.

**S1 File. Orthologous clusters associated with Fig 4a.** Orthologous clusters and singletons for the gene orthology analysis between the *D. pulicaria* genome, temperature-responsive genes in *D. pulex*, and kairomone-responsive genes in *D. pulex*. *D. pulex* genes were collected from the *Daphnia* Stressor Database. Data is in Excel format.

**S2 File. Orthologous clusters associated with Fig 4b.** Orthologous clusters and singletons for the gene orthology analysis between temperature-responsive *D. pulicaria* genes, kairomone-responsive *D. pulicaria* genes, temperature-responsive genes in *D. pulex*, and kairomone-responsive genes in *D. pulex*. *D. pulex* genes were collected from the *Daphnia* Stressor Database. Data is in Excel format.

**S3 File. Phenotype and life history data associated with Fig 1.**

## References

1. Carlson SM, Cunningham CJ, Westley PAH. Evolutionary rescue in a changing world. Trends in Ecology & Evolution. 2014;29(9):521–30.

2. Parmesan C. Ecological and Evolutionary Responses to Recent Climate Change. Annual Review of Ecology, Evolution, and Systematics. 2006;37(1):637–69.

3. Urban MC, Bocedi G, Hendry AP, Mihoub J-B, Pe’er G, Singer A, et al. Improving the forecast for biodiversity under climate change. Science [Internet]. 2016 Sep 9 [cited 2022 Jan 21]; Available from: https://www.science.org/doi/abs/10.1126/science.aad8466

4. Gienapp P, Teplitsky C, Alho JS, Mills J a., Merilä J. Climate change and evolution: Disentangling environmental and genetic responses. Molecular Ecology. 2008;17:167–78.

5. Merilä J, Hendry AP. Climate change, adaptation, and phenotypic plasticity: The problem and the evidence. Evolutionary Applications. 2014;7(1):1–14.

6. DeWitt TJ, Sih A, Wilson DS. Cost and limits of phenotypic plasticity. Trends in Ecology & Evolution. 1998;13(97):77–81.

7. Hendry AP, Farrugia TJ, Kinnison MT. Human influences on rates of phenotypic change in wild animal populations. Molecular Ecology. 2008;17(1):20–9.

8. Hoffmann AA, Shirriffs J, Scott M. Relative importance of plastic vs genetic factors in adaptive differentiation: Geographical variation for stress resistance in Drosophila melanogaster from eastern Australia. Functional Ecology. 2005;19(2):222–7.

9. Yampolsky LY, Zeng E, Lopez J, Williams PJ, Dick KB, Colbourne JK, et al. Functional genomics of acclimation and adaptation in response to thermal stress in Daphnia. BMC Genomics. 2014 Oct 4;15(1):859.

10. Whitehead A, Crawford DL. Neutral and adaptive variation in gene expression. 2007;103(May).

11. Aubin-Horth N, Renn SCP. Genomic reaction norms: using integrative biology to understand molecular mechanisms of phenotypic plasticity. Molecular ecology. 2009 Sep;18(18):3763–80.

12. Via S, Lande R. Genotype-Environment Interaction and the Evolution of Phenotypic Plasticity. Evolution. 1985;39(3):505–22.

13. Sørensen JG, Kristensen TN, Loeschcke V. The evolutionary and ecological role of heat shock proteins. Ecology Letters. 2003;6(11):1025–37.

14. Huang Y, Agrawal AF. Experimental Evolution of Gene Expression and Plasticity in Alternative Selective Regimes. PLOS Genetics. 2016;12(9):e1006336.

15. Kenkel C, Matz M V. Enhanced gene expression plasticity as a mechanism of adaptation to a variable environment in a reef-building coral. Nature Ecology and Evolution. 2016;1(3):059667.

16. Miner BE, De Meester L, Pfrender ME, Lampert W, Hairston NG. Linking genes to communities and ecosystems: Daphnia as an ecogenomic model. Proceedings of the Royal Society B: Biological Sciences. 2012;279(1735):1873–82.

17. Geerts AN, Vanoverbeke J, Vanschoenwinkel B, Van Doorslaer W, Feuchtmayr H, Atkinson D, et al. Rapid evolution of thermal tolerance in the water flea Daphnia. Nature Clim Change. 2015 Jul;5(7):665–8.

18. Williams PJ, Dick KB, Yampolsky LY. Heat tolerance, temperature acclimation, acute oxidative damage and canalization of haemoglobin expression in Daphnia. Evolutionary Ecology. 2012;26(3):591–609.

19. Yampolsky LY, Schaer TMM, Ebert D. Adaptive phenotypic plasticity and local adaptation for temperature tolerance in freshwater zooplankton. Proceedings Biological sciences / The Royal Society. 2014;281:20132744.

20. Stoks R, Govaert L, Pauwels K, Jansen B, De Meester L. Resurrecting complexity: The interplay of plasticity and rapid evolution in the multiple trait response to strong changes in predation pressure in the water flea Daphnia magna. Ecology Letters. 2016;19(2):180–90.

21. Kilham SS, Kreeger DA, Lynn SG, Goulden CE, Herrera L. COMBO: a defined freshwater culture medium for algae and zooplankton. Hydrobiologia. 1998;377(1):147–59.

22. Van Wagenen J, Miller TW, Hobbs S, Hook P, Crowe B, Huesemann M. Effects of Light and Temperature on Fatty Acid Production in Nannochloropsis Salina. Energies. 2012 Mar;5(3):731–40.

23. Coggins BL, Pearson AC, Yampolsky LY. Does geographic variation in thermal tolerance in Daphnia represent trade-offs or conditional neutrality? Journal of Thermal Biology. 2021 May 1;98:102934.

24. Glaholt SP, Kennedy ML, Turner E, Colbourne JK, Shaw JR. Thermal variation and factors influencing vertical migration behavior in Daphnia populations. J Therm Biol. 2016 Aug;60:70–8.

25. Palaima A, Spitze K. Is a jack-of-all-temperatures a master of none? An experimental test with Daphnia pulicaria (Crustacea: Cladocera). Evol Ecol Res. 2004;6(2):215–25.

26. Christjani M, Fink P, von Elert E. Phenotypic plasticity in three Daphnia genotypes in response to predator kairomone: evidence for an involvement of chitin deacetylases. Journal of Experimental Biology. 2016 Jun 1;219(11):1697–704.

27. Beauchamp DA. Seasonal and Diel Food Habits of Rainbow Trout Stocked as Juveniles in Lake Washington. Transactions of the American Fisheries Society. 1990 May 1;119(3):475–82.

28. Hembre LK. Effects of a rainbow trout stocking moratorium on the Daphnia species composition and water quality of Square Lake (Minnesota). Lake and Reservoir Management. 2019 Apr 3;35(2):127–39.

29. Knapp RA, Matthews KR, Sarnelle O. Resistance and Resilience of Alpine Lake Fauna to Fish Introductions. Ecological Monographs. 2001;71(3):401–21.

30. An H, Do TD, Jung G, Karagozlu MZ, Kim C-B. Comparative Transcriptome Analysis for Understanding Predator-Induced Polyphenism in the Water Flea Daphnia pulex. International Journal of Molecular Sciences. 2018 Jul;19(7):2110.

31. Zhang X, Blair D, Wolinska J, Ma X, Yang W, Hu W, et al. Genomic regions associated with adaptation to predation in Daphnia often include members of expanded gene families. Proceedings of the Royal Society B: Biological Sciences. 2021 Jul 28;288(1955):20210803.

32. Mitchell MD, Bairos-Novak KR, Ferrari MCO. Mechanisms underlying the control of responses to predator odours in aquatic prey. Journal of Experimental Biology. 2017 Jun 1;220(11):1937–46.

33. Roff DA. Evolutionary quantitative genetics. New York, NY: Chapman and Hall; 1997.

34. Core Team R. R: A language and environment for statistical computing. R Foundation for Statistical Computing. Vienna, Austria; 2018.

35. Kuznetsova A, Brockhoff PB, Christensen RHB. lmerTest package: tests in linear mixed effects models. Journal of Statistical Software. 2017;82:1–26.

36. Tukey JW. The Philosophy of Multiple Comparisons. Statistical Science. 1991 Feb;6(1):100–16.

37. Takele Assefa A, Vandesompele J, Thas O. On the utility of RNA sample pooling to optimize cost and statistical power in RNA sequencing experiments. BMC Genomics. 2020 Apr 19;21(1):312.

38. Chen S, Zhou Y, Chen Y, Gu J. fastp: an ultra-fast all-in-one FASTQ preprocessor. Bioinformatics. 2018 Sep 1;34(17):i884–90.

39. Wood DE, Lu J, Langmead B. Improved metagenomic analysis with Kraken 2. Genome Biology. 2019 Nov 28;20(1):257.

40. Sayers EW, Bolton EE, Brister JR, Canese K, Chan J, Comeau DC, et al. Database resources of the national center for biotechnology information. Nucleic Acids Res. 2021 Dec 1;gkab1112.

41. Grabherr MG, Haas BJ, Yassour M, Levin JZ, Thompson DA, Amit I, et al. Full-length transcriptome assembly from RNA-Seq data without a reference genome. Nat Biotechnol. 2011 Jul;29(7):644–52.

42. Manni M, Berkeley MR, Seppey M, Simão FA, Zdobnov EM. BUSCO Update: Novel and Streamlined Workflows along with Broader and Deeper Phylogenetic Coverage for Scoring of Eukaryotic, Prokaryotic, and Viral Genomes. Molecular Biology and Evolution. 2021 Oct 1;38(10):4647–54.

43. Jackson CE, Xu S, Ye Z, Pfrender ME, Lynch M, Colbourne JK, et al. Chromosomal rearrangements preserve adaptive divergence in ecological speciation [Internet]. 2021 Aug [cited 2021 Nov 2] p. 2021.08.20.457158. Available from: https://www.biorxiv.org/content/10.1101/2021.08.20.457158v1

44. Colbourne JK, Pfrender ME, Gilbert D, Thomas WK, Tucker A, Oakley TH, et al. The Ecoresponsive Genome of Daphnia pulex. Science. 2011 Feb 4;331(6017):555–61.

45. Nickel J, Schell T, Holtzem T, Thielsch A, Dennis SR, Schlick-Steiner BC, et al. Hybridization Dynamics and Extensive Introgression in the Daphnia longispina Species Complex: New Insights from a High-Quality Daphnia galeata Reference Genome. Genome Biology and Evolution. 2021 Dec 1;13(12):evab267.

46. Lee B-Y, Choi B-S, Kim M-S, Park JC, Jeong C-B, Han J, et al. The genome of the freshwater water flea Daphnia magna: A potential use for freshwater molecular ecotoxicology. Aquatic Toxicology. 2019 May 1;210:69–84.

47. Baldwin-Brown JG, Weeks SC, Long AD. A New Standard for Crustacean Genomes: The Highly Contiguous, Annotated Genome Assembly of the Clam Shrimp Eulimnadia texana Reveals HOX Gene Order and Identifies the Sex Chromosome. Genome Biology and Evolution. 2018 Jan 1;10(1):143–56.

48. Dobin A, Davis CA, Schlesinger F, Drenkow J, Zaleski C, Jha S, et al. STAR: ultrafast universal RNA-seq aligner. Bioinformatics. 2013 Jan;29(1):15–21.

49. Liao Y, Smyth GK, Shi W. featureCounts: an efficient general purpose program for assigning sequence reads to genomic features. Bioinformatics. 2014 Apr 1;30(7):923–30.

50. Love MI, Huber W, Anders S. Moderated estimation of fold change and dispersion for RNA-seq data with DESeq2. Genome biology. 2014;15(12):550.

51. Storey JD, Tibshirani R. Statistical significance for genomewide studies. PNAS. 2003 Aug 5;100(16):9440–5.

52. Chen H. VennDiagram: Generate High-Resolution Venn and Euler Plots [Internet]. 2021 [cited 2022 Jan 12]. Available from: https://CRAN.R-project.org/package=VennDiagram

53. Anders S, Huber W. Differential expression analysis for sequence count data. Nat Prec. 2010 Mar 15;1–1.

54. Anderson MJ. Permutational Multivariate Analysis of Variance (PERMANOVA). In: Wiley StatsRef: Statistics Reference Online [Internet]. John Wiley & Sons, Ltd; 2017 [cited 2022 Feb 8]. p. 1–15. Available from: https://onlinelibrary.wiley.com/doi/abs/10.1002/9781118445112.stat07841

55. Oksanen J, Blanchet FG, Friendly M, Kindt R, Legendre P, McGlinn D, et al. vegan: Community Ecology Package [Internet]. 2020. Available from: https://CRAN.R-project.org/package=vegan

56. Gene Ontology Consortium. The Gene Ontology resource: enriching a GOld mine. Nucleic Acids Res. 2021 Jan 8;49(D1):D325–34.

57. Young MD, Wakefield MJ, Smyth GK, Oshlack A. Gene ontology analysis for RNA-seq: accounting for selection bias. Genome biology. 2010;11(2):R14.

58. Benjamini Y, Hochberg Y. Controlling the False Discovery Rate: A Practical and Powerful Approach to Multiple Testing. Journal of the Royal Statistical Society: Series B (Methodological). 1995 Jan 1;57(1):289–300.

59. Ravindran SP, Lüneburg J, Gottschlich L, Tams V, Cordellier M. Daphnia stressor database: Taking advantage of a decade of Daphnia ‘-omics’ data for gene annotation. Sci Rep. 2019 Jul 31;9(1):11135.

60. Colbourne JK, Singan VR, Gilbert DG. wFleaBase: the Daphnia genome database. BMC Bioinformatics. 2005 Mar 7;6:45.

61. Xu L, Dong Z, Fang L, Luo Y, Wei Z, Guo H, et al. OrthoVenn2: a web server for whole-genome comparison and annotation of orthologous clusters across multiple species. Nucleic Acids Res. 2019 Jul 2;47(W1):W52–8.

62. Stibor H. Predator induced life-history shifts in a freshwater cladoceran. Oecologia. 1992;92(2):162–5.

63. Riessen HP. Predator-induced life history shifts in Daphnia: a synthesis of studies using meta-analysis. Canadian Journal of Fisheries and Aquatic Sciences. 1999;56:2487–94.

64. Tseng M, O’Connor MI. Predators modify the evolutionary response of prey to temperature change. Biology letters. 2015;11(12):20150798.

65. Zhang C, Jansen M, De Meester L, Stoks R. Energy storage and fecundity explain deviations from ecological stoichiometry predictions under global warming and size-selective predation. The Journal of animal ecology. 2016 Nov;85(6):1431–41.

66. Luhring TM, DeLong JP. Predation changes the shape of thermal performance curves for population growth rate. Current zoology. 2016 Oct;62(5):501–5.

67. Luhring TM, Vavra JM, Cressler CE, DeLong JP. Predators modify the temperature dependence of life-history trade-offs. Ecology and Evolution. 2018 Sep 1;8(17):8818–30.

68. Riessen HP. Water temperature alters predation risk and the adaptive landscape of induced defenses in plankton communities. Limnology and Oceanography. 2015 Nov 1;60(6):2037–47.

69. Miller LP, Matassa CM, Trussell GC. Climate change enhances the negative effects of predation risk on an intermediate consumer. Global Change Biology. 2014;20:3834–3844.

70. von Elert E, Agrawal MK, Gebauer C, Jaensch H, Bauer U, Zitt A. Protease activity in gut of Daphnia magna: evidence for trypsin and chymotrypsin enzymes. Comparative Biochemistry and Physiology Part B: Biochemistry and Molecular Biology. 2004;137(3):287–96.

71. Colbourne JK, Eads BD, Shaw J, Bohuski E, Bauer DJ, Andrews J. Sampling Daphnia’s expressed genes: preservation, expansion and invention of crustacean genes with reference to insect genomes. BMC Genomics. 2007;8(1):217.

72. Williamson CE, Fischer JM, Bollens SM, Overholt EP, Breckenridge JK. Toward a more comprehensive theory of zooplankton diel vertical migration: Integrating ultraviolet radiation and water transparency into the biotic paradigm. Limnology and Oceanography. 2011;56(5):1603–23.

73. Tams V, Nickel JH, Ehring A, Cordellier M. Insights into the genetic basis of predator-induced response in Daphnia galeata. Ecology and Evolution. 2020;10(23):13095–108.

74. Orsini L, Brown JB, Shams Solari O, Li D, He S, Podicheti R, et al. Early transcriptional response pathways in Daphnia magna are coordinated in networks of crustacean-specific genes. Molecular Ecology. 2018;27(4):886–97.

75. Hahn MA, Effertz C, Bigler L, von Elert E. 5α-cyprinol sulfate, a bile salt from fish, induces diel vertical migration in Daphnia. Pohnert G, Baldwin IT, Pohnert G, editors. eLife. 2019 May 2;8:e44791.

76. Loose CJ, Elert E, Dawidowicz P. Chemically-induced diel vertical migration in Daphnia: a new bioassay for kairomones exuded by fish. Archiv für Hydrobiologie. 1993;329–37.

77. Diel P, Kiene M, Martin-Creuzburg D, Laforsch C. Knowing the Enemy: Inducible Defences in Freshwater Zooplankton. Diversity. 2020 Apr;12(4):147.

78. Rozenberg A, Parida M, Leese F, Weiss LC, Tollrian R, Manak JR. Transcriptional profiling of predator-induced phenotypic plasticity in Daphnia pulex. Frontiers in Zoology. 2015 Jul 25;12(1):18.

79. Miehls ALJ, McAdam AG, Bourdeau PE, Peacor SD. Plastic response to a proxy cue of predation risk when direct cues are unreliable. Ecology. 2013;94(10):2237– 48.

80. Riessen HP, Gilbert JJ. Divergent developmental patterns of induced morphological defenses in rotifers and Daphnia: Ecological and evolutionary context. Limnology and Oceanography. 2019;64(2):541–57.

81. Suppa A, Caleffi S, Gorbi G, Marková S, Kotlík P, Rossi V. Environmental conditions as proximate cues of predation risk inducing defensive response in Daphnia pulex. Biologia. 2021 Feb 1;76(2):623–32.

82. Lampert W. The Adaptive Significance of Diel Vertical Migration of Zooplankton. Functional Ecology. 1989;3(1):21–7.

83. Stich H-B, Lampert W. Growth and reproduction of migrating and non-migrating Daphnia species under simulated food and temperature conditions of diurnal vertical migration. Oecologia. 1984 Feb;61(2):192–6.

84. Hembre LK, Peterson LA. Evolution of predator avoidance in a Daphnia population: evidence from the egg bank. Hydrobiologia. 2013 Jan 1;700(1):245–55.

85. Ślusarczyk M. Predator-induced diapause in Daphnia magna may require two chemical cues. Oecologia. 1999 May 1;119(2):159–65.

86. Stabell OB, Ogbebo F, Primicerio R. Inducible Defences in Daphnia Depend on Latent Alarm Signals from Conspecific Prey Activated in Predators. Chemical Senses. 2003 Feb 1;28(2):141–53.

87. Schoeppner NM, Relyea RA. Interpreting the smells of predation: how alarm cues and kairomones induce different prey defences. Functional Ecology. 2009;23(6):1114–21.

88. >Ślusarczyk M, Rygielska E. Fish Faeces as the Primary Source of Chemical Cues Inducing Fish Avoidance Diapause in Daphnia Magna. Hydrobiologia. 2004 Sep 1;526(1):231–4.

89. Boeing WJ, Ramcharan CW, Riessen HP. Multiple predator defence strategies in Daphnia pulex and their relation to native habitat. Journal of Plankton Research. 2006 Jun 1;28(6):571–84.

90. Nagano M, Doi H. Ecological and evolutionary factors of intraspecific variation in inducible defenses: Insights gained from Daphnia experiments. Ecology and Evolution. 2020;10(16):8554–62.

91. Lima TG, Willett CS. Locally adapted populations of a copepod can evolve different gene expression patterns under the same environmental pressures. Ecol Evol. 2017 May 9;7(12):4312–25.

92. Roy Chowdhury P, Frisch D, Becker D, Lopez JA, Weider LJ, Colbourne JK, et al. Differential transcriptomic responses of ancient and modern Daphnia genotypes to phosphorus supply. Mol Ecol. 2015 Jan;24(1):123–35.

